# Effects of Deficient Glycosylation and Deglycosylation on Sperm Condition in Zebrafish (*Danio rerio*)

**DOI:** 10.64898/2026.07.01.735899

**Authors:** Kelly McGraw, Marie Mooney

**Affiliations:** Department of Biology, University of North Florida, Jacksonville, FL 32224

## Abstract

Congenital disorders of glycosylation and deglycosylation are rare, serious, and lethal disorders afflicting humans. CDGs and CDDGs result in loss of function enzymes which fail to build or break down oligosaccharides on proteins. This can produce protein aggregates and, in turn, reactive oxygen species that harm the cell eventually leading to autophagy and apoptosis. Because sperm contain high concentrations of polyunsaturated fatty acids, they are especially sensitive to these effects, which is understood as one of the leading factors in human male infertility. Sperm are developed in zebrafish similarly to humans and are useful models to examine human reproductive health, as well as genetic disorders. The combination of these advantages makes the analysis of sperm from zebrafish with heterozygous ALG1 or DPAGT1 CDGs or the NGLY1 CDDG suitable. Analysis of sperm concentration, motility, status, viability, and hypoosmotic swelling demonstrated the effects of these disorders on sperm quality. Results showed a significant decrease in sperm concentration, motility, and hypoosmotic swelling for all mutant zebrafish compared to the wild type. This suggests that CDGs and CDDGs influence the amount of sperm produced, the percentage of sperm cells that are mobile, and the integrity of the plasma membrane.

## Introduction

The small tropical freshwater zebrafish, *Danio rerio*, are a commonly used animal model in modern research for their easy care, maintenance and developmental advantages such as their transparent larvae. The zebrafish and human genomes retain 70% of the same genes and are useful models to examine human genetic disorders (Howe et al., 2013). In addition to the similar genome, both humans and zebrafish are closely related in their production and regulation of gametes, and tubule organization of Sertoli cells for spermatogenesis (Hoo et al., 2016). Because of this similarity in sperm development, the sperm produced by zebrafish can reveal how genetic disorders may affect the quality or condition of mature sperm.

In early spermatogenesis, spermatocytes include the nucleus, the Golgi complex, and endoplasmic reticulum (ER) associated with the nuclear envelope. As the spermatocytes undergo meiosis, the ER disassociates with each division until it is exocytosed as the spermatid matures (Nistal et al., 2008). Development of spermatocytes into mature spermatozoa is dependent on the number of Sertoli cells as they regulate crucial signaling in the developmental process in the testis. The number of Sertoli cells is the largest limiting factor of spermatogenesis. Their signaling organizes the growth factors for sperm development at each stage, thereby allowing for fewer instances of spermatozoa apoptosis, which the Sertoli cells are responsible for phagocytizing (Schulz et al., 2010). The phagocytosis of Sertoli cells in fish is highly efficient, clearing residual bodies from the developing sperm quickly and recycling the material.

According to Schulz et al. (2010) of all the spermatocytes that are destined to differentiate into spermatozoa, 30-40% could theoretically become apoptotic, yet histological sections show only 10% of spermatocytes are apoptotic, demonstrating the efficiency of the Sertoli cells. Because Sertoli cells are vital to the development and survival of spermatocytes maturing into spermatozoa, irregularities in their cellular communication and functional efficiency are reflected in the condition of sperm cells.

Congenital disorders of glycosylation (CDGs) are rare and serious human disorders that affect the synthesis of oligosaccharides to proteins or lipids, which are key communicators in cell-cell signaling. One such CDG is the ALG1-CDG which results from a mutation in the ALG1 enzyme, mannosyltransferase-1, embedded in the ER membrane that normally catalyzes the first mannosylation of N-acetylglucosamine (GlcNAc). Mutations in ALG1 that cause loss of enzymatic function result in shortened oligosaccharides (Schwarz et al., 2004). Loss-of-function mutations down-regulate proteins related to cell growth, cell survival, and decrease proteins associated with mitochondria. Additionally, an increase in autophagy is found in mutant cells which suggest possible mitochondrial and cellular stress or damage (Budhraja et al., 2024). The ultimate result of the ALG1-CDG is death during adolescence. Likewise, in the model organism, zebrafish, the homozygous mutant fish are lethal several days post fertilization. To this date, there are no known cures or treatments for this disorder which makes the need for further research on understanding this disorder necessary.

A similar CDG to ALG1 is DPAGT1, an enzyme in the same glycosylation pathway, positioned ahead of ALG1. DPAGT1 encodes the enzyme N-acetylglucosamine-1-phosphotransferase which catalyzes the first step in N-glycosylation by adding phosphate and GlcNAc to dolichol phosphate (Hyde et al., 2022). Loss-of-function mutations in DPAGT1 have been shown to increase both ER stress and apoptosis. Both findings are linked due to the ER inducing apoptosis caused by the stress of protein accumulation and aggregation (Hyde et al., 2022). Like the ALG1-CDG, the DPAGT1-CDG ultimately causes death in adolescence, and the zebrafish animal model cannot survive the homozygous mutation a few days post fertilization.

The promotion of autophagy, and eventually apoptosis in CDGs, may be linked to the cellular stress caused by reactive oxygen species (ROS) produced by the aggregation of misfolded, shortened glycoproteins. Normally, the cell uses the unfolded protein response (UPR) to handle the misfolding of proteins; however, the mutated glycoprotein cannot be folded into its correct conformation, so it is sent for degradation (Daniel et al., 2025). The endoplasmic-reticulum-associated degradation (ERAD) system is then responsible for the transport of the misfolded proteins out of the ER. The misfolded protein is finally degraded in the cytoplasm by ubiquitin tagging or autophagy (Li et al., 2022). It is well understood that autophagy switches to apoptosis when cellular stress becomes too great; therefore, the large increase in mitochondrial stress, oxidative stress, and ER stress likely influences the cell to become apoptotic when mutated by a CDG.

In contrast to congenital disorders of glycosylation, there is another serious, rare disorder involved in the degradation of glycans, congenital disorders of deglycosylation (CDDGs). The most prevalent CDDG is the mutation of the NGLY1 enzyme, N-glycanase 1, which is responsible for the non-lysosomal degradation of misfolded proteins within the ERAD system (Shyr et al., 2025). NGLY1 deficiency causes the upregulation of ERAD and an increase in lipid peroxidation, heightening the levels of ROS within the cell. Large amounts of radical hydroxide can cause damage to organelles and the plasma membrane, decreasing its integrity. As a result, autophagy is increased by these stress-induced pressures (Shyr et al., 2025). Like the CDGs, NGLY1-CDDG causes death in adolescence and in most animal models. The zebrafish can survive the homozygous loss of function in NGLY1, but for comparison, only the heterozygous mutant of this disorder is analyzed.

The effects of these CDGs and CDDGs on sperm have not yet been studied. Cellular stress can damage and reduce the quality of sperm cells, which establishes the basis of this study. Chronic stress in zebrafish dysregulates the production of spermatozoa, altering the concentration and motility of the mature cell, and downregulating the pathway which eliminates missense mutation in RNA (Valcarce et al., 2023). Here, the effectors of chronic stress being tested are ALG1, DPAGT1, and NGLY1 heterozygous mutant zebrafish, respectively. As previously described, CDGs and CDDGs increase oxidative, ER, and mitochondrial stress due to the accumulation of protein aggregates and excessive ROS. The presence of the ER in early spermatocytes, along with the mitochondria that remain in the sperm tail and are crucial for sperm function, creates the potential for harmful effects due to these disorders (Boryshpolets et al., 2009). Additionally, sperm cells are highly sensitive to ROS due to their high content of polyunsaturated fatty acids which are vulnerable to lipid peroxidation (Asadi et al., 2021).

Because of this, a heterozygous mutant is expected to have higher levels of aggregation and ROS compared to the wild type; therefore, heterozygous mutants are expected to produce lower quality sperm. There is also the likelihood that the supporting somatic Sertoli cells are impacted by these disorders which in turn causes or amplifies the risk of lower sperm quality. However, it should be noted that despite these hypothesized effects, heterozygous mutant fish are haplosufficient and continue to produce progeny that follows Mendelian ratios.

To analyze the impacts of loss-of-function mutations in CDG/CDDG enzymes on the quality of sperm, we assess 1) sperm concentration, 2) sperm motility and status, 3) sperm viability, and 4) hypoosmotic swelling. General analysis of sperm concentration provides insight to the possible dysregulation in sperm production (Valcarce et al., 2023). The motility or percentage of mobile spermatozoa demonstrates the effect of mitochondrial stress as well as its relation to viability. For deeper insights on mitochondrial condition, a status or objective sperm speed analysis can signify the amount of energy (ATP) a spermatozoon has (Boryshpolets et al., 2009). Viability analyses have been used for numerous sperm quality studies, and here it can demonstrate the ability of a mutant cell, spermatozoon or Sertoli, to survive when impacted by stress (Cattelan and Gasparini, 2021). Finally, a hypoosmotic swelling test (HOST) is one of the most desirable parameters to test against CDGs and CDDGs because of the effects oxidative stress has on membrane integrity. If spermatozoa are damaged by oxidative stress, the HOST would result in cellular death and eventually lyse (Cabrita et al., 1999). Because of oxidative, ER, and mitochondrial stress caused by the protein aggregation because of ALG1, DPAGT1, and NGLY1 mutations, it is expected that concentration, motility, status, viability, and hypoosmotic swelling of zebrafish spermatozoa will be decreased significantly compared to the wild type.

## Materials and Methods

### Zebrafish Husbandry

Heterozygous mutant genotypes were previously created from WT [AB] lines using CRISPR-Cas9 technology at Duke University or Lurie Children’s Hospital/Northwestern University under their IACUC protocols. All zebrafish in this study were bred and raised in the University of North Florida zebrafish housing facility and were maintained at standard conditions on a recirculating water system with room temperature set at 28.5°C and a diurnal light cycle. All fish used were approximately 18 months old and were fed using Gemma 500 every afternoon. All research activities were approved by the University of North Florida Institutional Animal Care and Use Committee, Protocol IA 21-002.

### Sperm Stripping

In the evening prior to sperm collection, three males and one female of the same genotype were placed in breeding tanks with males and the female separated with a clear divider. Tanks were left overnight off system on a breeding rack. Sperm collection occurred in the morning with low lighting. Male fish were anesthetized with 0.2 mg/mL MS-222 until operculum movement halted or slowed significantly and fish turned belly-up. Fish were removed and rinsed in an isotonic solution before being blotted dry with a paper towel. Then, they were placed on a sponge dampened with system water and the urogenital pore was blotted dry to minimize spontaneous sperm activation (Cheng et al., 2021). Gentile abdominal pressure was applied to the fish, and sperm was collected using a microcapillary tube. Each sperm sample was equalized to 5 µL in a sperm extender. Collection was performed 4-5 days apart to allow for recovery, stress minimization, and for the most adequate time for spermatogenesis (Cattelan and Gasparini, 2021).

### Concentration of Sperm per Fish

Each sperm collection was added to a 96-well plate and diluted 1:6 with 1x Hank’s Balanced Salt Solution (HBSS). Approximately 10 wells were filled with an equal volume of HBSS as a blank. The plate was read with a spectrophotometer at 400 nm (A400). Using the absorbance values and the equation for sperm concentration produced by Matthews et al. (2023), the concentration of sperm per fish was calculated and averaged (Equation 1). These values were then divided by 1 million for simplicity.

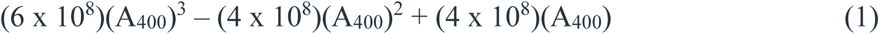

### Extender Comparison

To eliminate the possibility of extender influence on sperm analysis, a comparison of extenders was made. The three extenders used were made from 10x HBSS. The solution was diluted to 1x, creating a HBSS solution with a pH of 5.5. The second extender used 1x HBSS and added sodium bicarbonate until the pH reached 7.2. The third extender used 1x HBSS pH 7.2 and added 15 mg/mL of bovine serum albumin (Cheng et al., 2021) to create a Modified Hank’s Balanced Salt Solution (MHBSS).

Sperm were stripped from nine WT [AB] zebrafish with three fish pooled for each extender. Pooled sperm was normalized in one of the three extenders then diluted 1:5 into the same extender. To keep sperm for an extended time, samples were kept on ice. Aliquots of sperm were activated with DI water and examined for motility every 15 minutes for 120 minutes. Three measurements were taken for each sample every time.

Results of the motility examination were averaged per time point and extender before being plotted (Figure 2). The best performing extender was then utilized for the motility, status, viability, and HOST.

**Figure 1.**
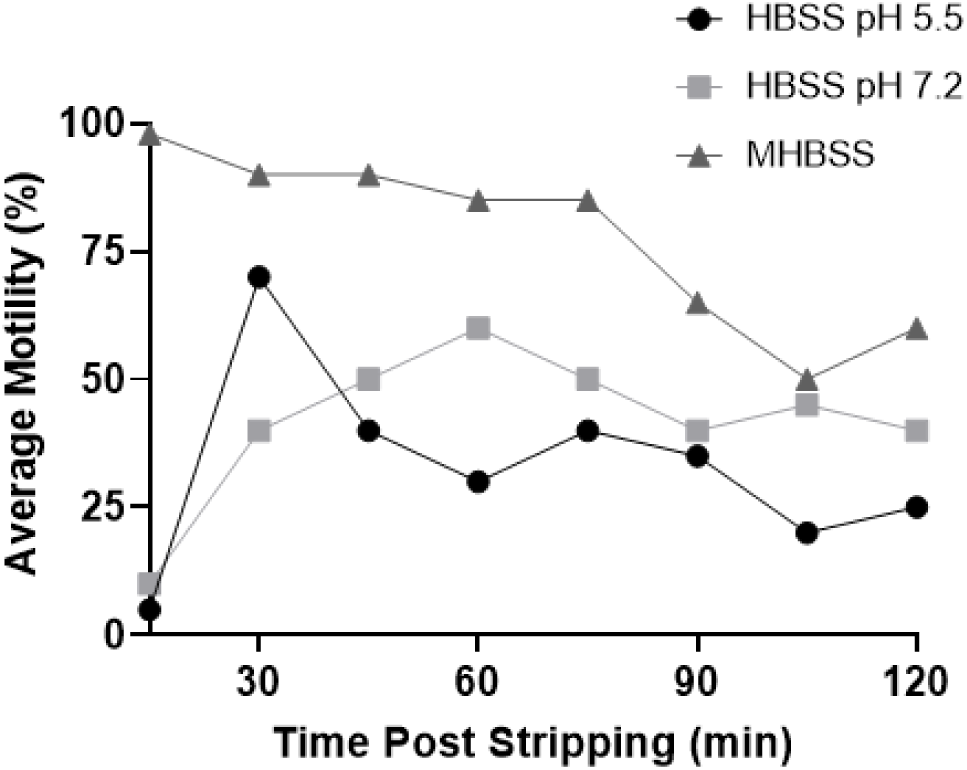
Comparison of Sperm Extenders on Motility Over Time. Sperm was stripped from WT [AB] fish (n = 3 per extender) and pooled according to extender group. Percent motility was measured every 15 minutes for 2 hours. Turkey’s multiple comparison test found MHBSS was significantly different from HBSS pH 5.5 and HBSS pH 7.2 (p<0.0001 and p<0.01, respectively).

**Figure 2.**
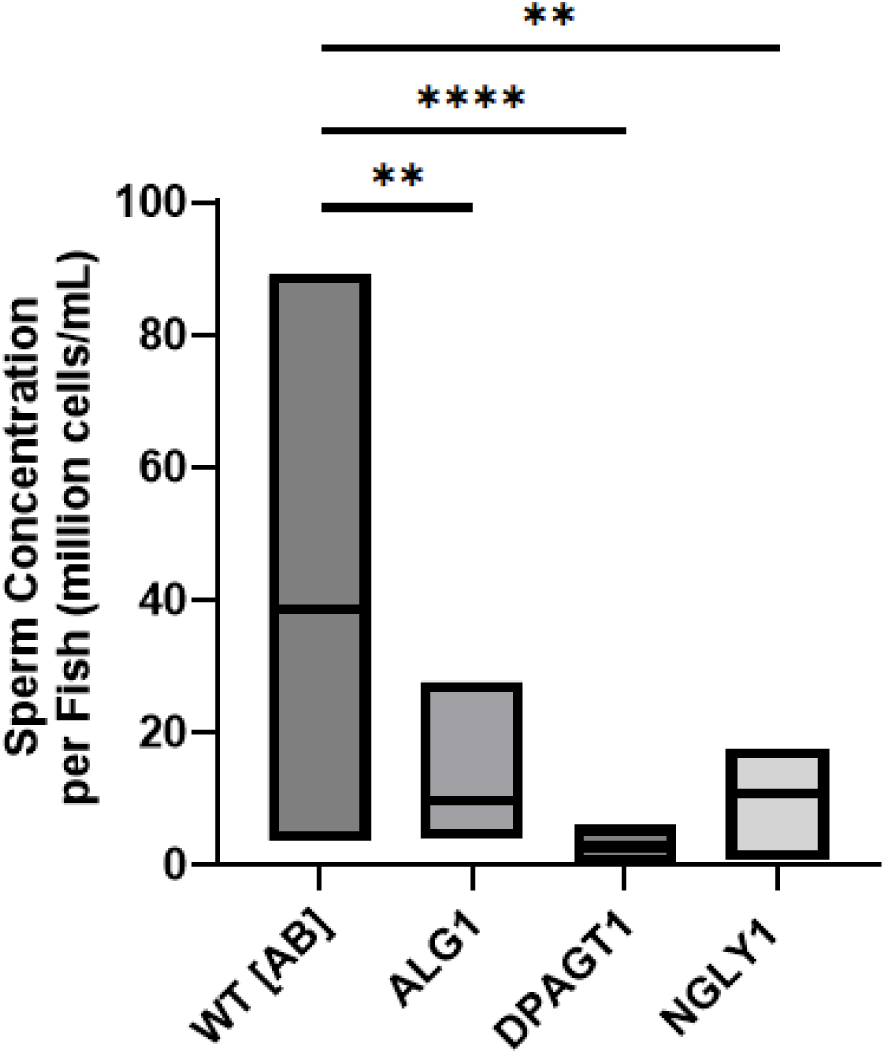
Comparison of Stripped Sperm Concentration per Fish. Sperm was collected and immediately added to a 1x solution of HBSS in a 96-well plate. Each sample was diluted 1:6 with additional HBSS and their absorbance was read at 400nm. Using the concentration equation (Equation 1), absorbance was converted into cells/mL. Turkey’s multiple comparison test found ALG1, DPAGT1, and NGLY1 were significantly different than WT [AB] (p<0.01).

### Motility and Status

Three fish per genotype were stripped, equalized in MHBSS, and pooled together. Additional MHBSS was added to make a final 1:5 dilution. All samples were kept on ice to extend working time. Aliquots of sperm were placed on slides warmed to 38°C, and an equal amount of warm DI water was added to activate the sperm. Motility was determined by examining five fields of view and averaging the percent motility of each field. Similarly, status was determined by examining five fields of view and averaging the ranking for each field. Status was ranked 1 to 4 with 1 being the slowest moving sperm and 4 being the fastest moving sperm.

### Viability

Three fish per genotype were stripped, equalized in MHBSS, and pooled together. Additional MHBSS was added to make a final 1:2 dilution. For each sample, an equal amount of eosin-nigrosin sperm viability stain solution was added for a final 1:2 sperm-stain dilution. The sperm-stain solution was smeared onto a slide warmed to 38°C and allowed to completely air dry. Eosin-stained dead, nonviable cells pink and viable; live cells remain white (Figure 4B, 4D). Nigrosin acted as a negative stain for better visualization of the primary eosin stain. Cells were counted with different fields of view until cell counts were equal to or greater than 300.

**Figure 3.**
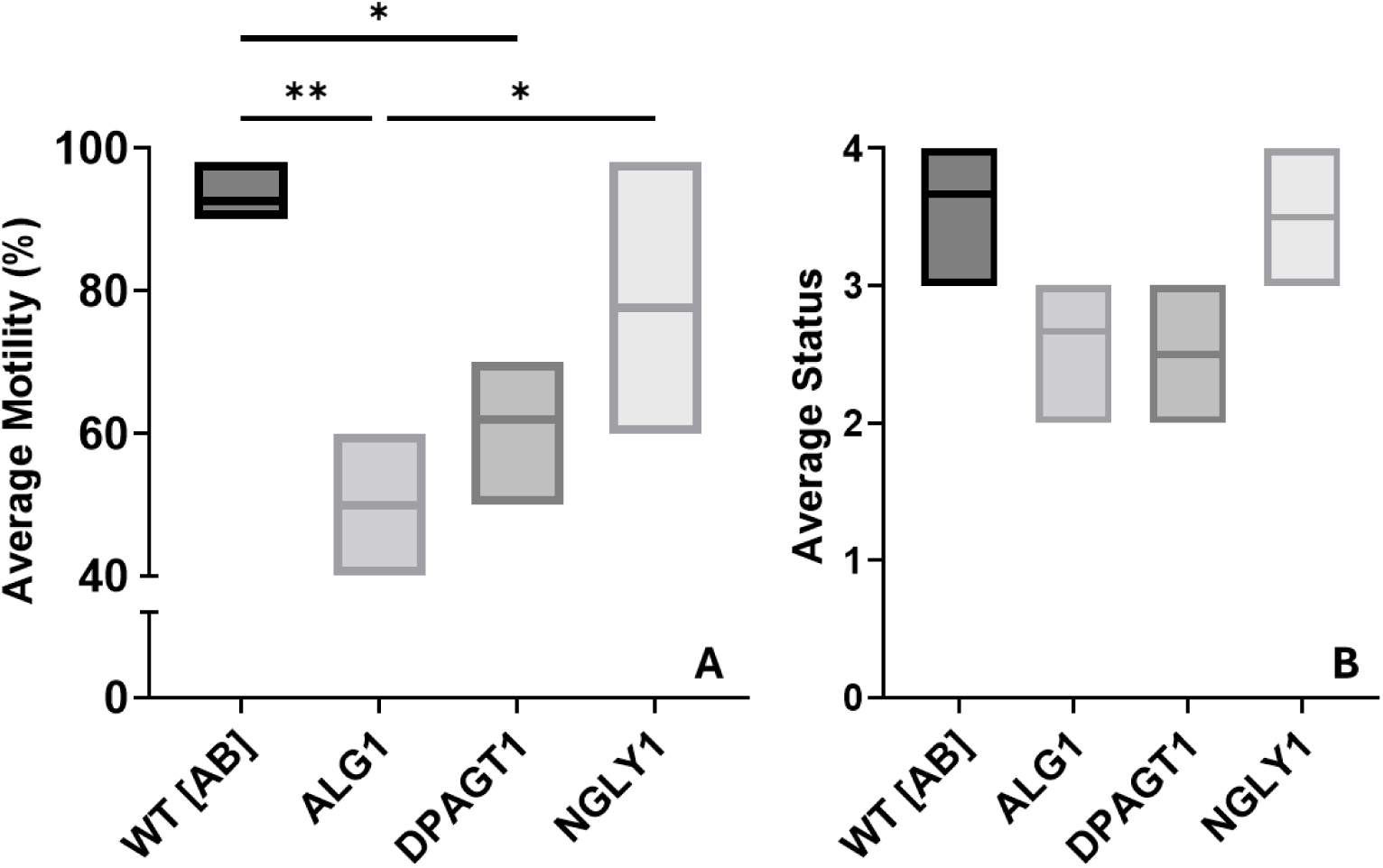
Comparison of Sperm Status and Motility to Genotype. Sperm were stripped from fish of each genotype (n = 3 per genotype) and placed into MHBSS. A 5 µL aliquot of sperm activated with equal part DI water. Motility and status were measured under a microscope at high magnification. Measurements were repeated three times per genotype. A) Motility of sperm was measured objectively, and measurements were averaged with three replicates each. Significance was determined by Turkey’s multiple comparison test. B) Status of sperm movement was ranked objectively 1 through 4 and measurements were averaged per genotype. Significance was determined by Turkey’s multiple comparison test.

**Figure 4.**
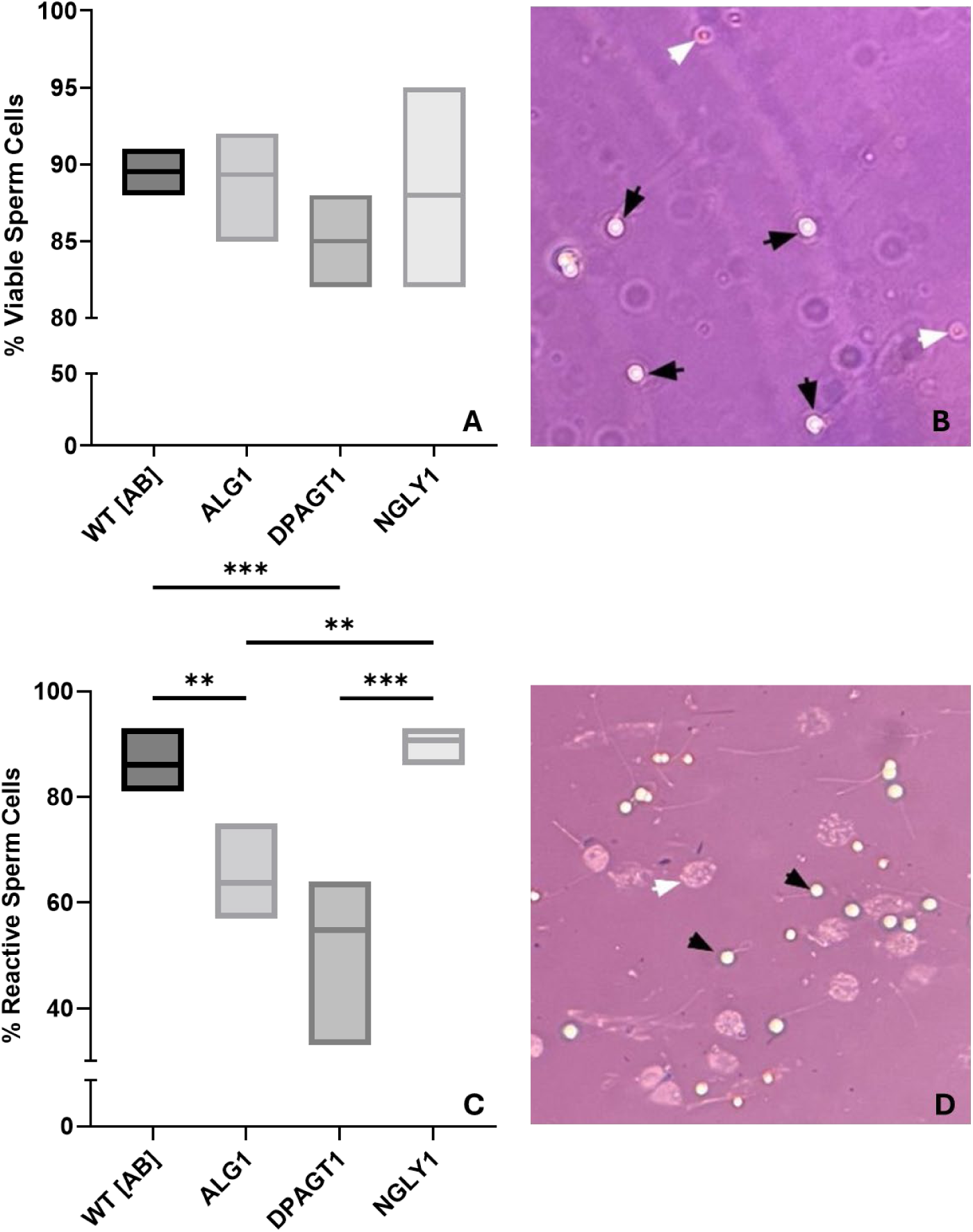
Comparison of Sperm Cell Viability and HOST Reactivity to Genotype. Sperm were stripped from fish of each genotype (n = 3 per genotype) and placed into MHBSS for a 1:2 dilution. A) Sperm in solution were mixed with an equal part eosin-nigrosin staining solution. Sperm-stain solution was smeared on a slide before being measured under a microscope. Cells were counted until n > 300. Significance was determined by Turkey’s multiple comparison test. B) Viability stain of DPAGT1 sperm cells at 400x total magnification. Black arrows point to white, viable cells. White arrows point to pink, dead cells. C) Sperm was added to an equal part of DI water and left in a 38°C water bath for 30 minutes. Then, an equal part of eosin-nigrosin staining solution was added to the sperm-stain solution and was smeared onto a slide. Significance was determined by Turkey’s multiple comparison test. D) HOST stain of DPAGT1 sperm cells at 400x total magnification. Black arrows point to reactive, swelling cells. The white arrow points to unreactive, lysed cells.

### Hypoosmotic Swelling Test

HOST was performed by using pooled, equalized sperm with three fish per genotype added to an equal amount of DI water creating a 1:2 dilution. This solution was left in a 38°C water bath for 30 minutes. Then, the HOST sperm solution was added to an equal part of the eosin-nigrosin stain for a final 1:2 HOST-stain dilution. The final solution was smeared on a warmed slide and allowed to air dry completely. Reactive, swollen cells appear white, coiled, or with droplets on the sperm tail. Unreactive cells appear either lysed or with straight, unswollen tails (Cabrita et al., 1999). Cells were counted with different fields of view until cell counts were equal to or greater than 300.

### Statistical Analyses

Each analysis was analyzed using an ANOVA one-way test on the GraphPad Prism 10. Additionally, each parameter was analyzed with Tukey’s multiple comparisons, Brown-Forsythe test, and Barlett’s test. Data was shown graphically with mean ± SEM (\**p*<0.05; \*\**p*<0.01; \*\*\**p*<0.001; ns: no significant differences).

## Results

To examine the effects of CDGs and CDDGs on sperm condition, a series of four analyses were performed: quantification of sperm production, analysis of sperm movement via motility and status measures, determination of viability by measuring the proportion of living cells, and analysis of membrane integrity of sperm via the HOST. Overall results demonstrate lower concentration, motility, and swelling reactivity in mutant genotypes.

### Analysis of Sperm Extenders

Three extenders analyzed included HBSS pH 5.5, HBSS pH 7.2, and MHBSS. Motility of sperm overtime in each extender was recorded using WT [AB] fish and plotted in Figure 2. No significant difference was found between HBSS pH 5.5 and 7.2 (*p* = 0.5664). Motility of sperm in MHBSS differed significantly compared to HBSS pH 5.5 ( *p*< 0.0001) and HBSS pH 7.2 (*p* = 0.001). Brown-Forsythe and Bartlett’s tests found no significant difference of SD (*p=*0.7878 and p=0.8071 respectively). For the following sperm analyses, MHBSS was chosen as the extender.

### Concentration Analysis

Absorbance values were used to calculate the concentration of sperm from equation 1. Calculation of sperm concentration per fish per genotype produced significantly different results for ALG1, DPAGT1, and NGLY1 compared to WT [AB] (*p* = 0.0045, <0.0001, and 0.0022 respectively). No significance was found between each of the three mutant genotypes with each other (Figure 1). Brown-Forsythe and Bartlett’s tests found SD to be significantly different (*p*<0.0001 and *p*<0.0001 respectively).

### Sperm Motility and Status

The motility of WT [AB] was found to be significantly different from two of the mutant genotypes with the greatest difference observed with ALG1 (*p*=0.0027). WT [AB] differed with DPAGT1 (*p* = 0.0131) but did not significantly differ from NGLY1 (*p* = 0.3051). Additionally, ALG1 was significantly different than the results of NGLY1 motility (*p* = 0. 0253) (Figure 3A). The status, or objective speed, was not found to be significantly different between WT [AB] and any of the three mutant genotypes (Figure 3B). The Brown-Forsythe test found no significant difference in the SD of motility or status (*p*=0.5213 and *p*=0.8678 respectively).

### Sperm Viability and Swelling Reactivity

Analysis of viable cell counts from eosin-nigrosin staining showed no significant difference between WT [AB] viability compared to the viability of the three mutant lines individually (*p*>0.05). Figure 4A showed a trend toward higher viability in NGLY1 sperm compared to WT [AB] sperm cells and a trend toward lower viability in DPAGT1 sperm compared to WT [AB] sperm cells. A representation of viable and nonviable sperm cells was shown in Figure 4B. The Brown-Forsythe test showed no significant difference in the SD of viability *(p*=0.7487).

The HOST test found a significant difference in the reactivity of two mutant sperm genotypes in hypoosmotic conditions compared to the WT [AB] sperm. WT [AB] differed the most to DPAGT1 (*p* = 0.0002), then ALG1 (*p* = 0.0037), and did not differ with NGLY1 (*p* = 0.8897). The reactivity of the mutants was found to significantly differ between NGLY1 and ALG1 (*p*=0.0016) and DPAGT1 (*p*=0.0001) respectively (Figure 4C). A representation of reactive cells and unreactive cells was shown in Figure 4D. The Brown-Forsythe test showed no significant difference in the SD of HOST (*p*=0.5717).

## Discussion

With consideration of CDGs and CDDG impacts on intracellular integrity, the examination of sperm revealed crucial insight into the effects these disorders had on sperm conditions. It was hypothesized that cellular stress caused apoptosis and decreased membrane integrity which were to be reflected in sperm concentration, motility, status, viability, and hypoosmotic swelling. Analysis of these parameters confirmed concentration, motility, and hypoosmotic swelling were significantly decreased compared to wild type sperm. Status of sperm did not differ from wild type for any of the three mutant genotypes (Figure 3A). These findings were indicative of a negative sub-cellular condition that caused the decrease in sperm quality, which was likely the result of the CDG or CDDG mutation. Future research should focus on the quantification of ROS, Sertoli cells, and mitochondria to support these findings.

The concentration per zebrafish from each genotype revealed a significant decrease in sperm concentration for mutant fish compared to the WT [AB] line. DPAGT1 demonstrated the greatest decrease in sperm concentration. This dramatic decrease in concentration may be attributed to the ER and oxidative stress caused by the ROS produced from protein aggregates. However, in a study that compared chronic stress to the concentration of zebrafish sperm, they had found no change in concentration (Valcarce et al., 2021). It should be noted that the stress factors in their study were environmental not genetic. In a study of human sperm impacted by ROS, concentration of sperm cells was significantly lower linking male infertility to ROS levels (Agarwal et al., 2006). With these prior studies in mind, cellular stress caused by ROS and not an environmental factor may have caused a decrease in sperm concentration for mutant genotypes. Further research quantifying the levels of ROS in the sperm would be necessary to confirm this hypothesis.

Motility was another parameter that was affected significantly by the presence of a CDG and not a CDDG. DPAGT1 and ALG1 mutants were both found to be less motile than WT [AB]. The motility of ALG1 mutant sperm decreased the greatest, differing significantly from both WT [AB] and NGLY1 mutants (Figure 3A). In both the studies done by Valcarce et al. (2021) and Agrarwal et al. (2006), motility was decreased in the presence of stressors or the presence of ROS. Unexpectedly, the status of sperm did not change significantly for any of the three mutants compared to WT [AB], reflecting the possibility that mitochondrial activity and ATP availability were not impaired by their loss of function. Therefore, motility was highly affected by the presence of CDGs, especially in ALG1 mutants. Status of mutant sperm were not affected by the presence of CDGs or CDDG. To further support these findings, the use of an ATP colorimetric assay could be used to quantify the amount of energy produced by sperm mitochondria. Motility was also affected significantly by the choice of sperm extender. Sperm motility dramatically decreased when stored in HBSS pH 5.5 and pH 7.2. Motility between these two groups did not significantly differ, though pH 7.2 may perform slightly better. The MHBSS performed the best out of the three conditions with the highest motility of 98% at 15 minutes (Figure 2). A similar result was found by Cheng et al. (2021) when testing MHBSS against standard extenders E400 and Kurokura. They demonstrated MHBSS’s ability to nearly replicate the motility of WT [AB] sperm to the standard extenders over a period of 24 hours. As the time frame used in their experiment was beyond the scope for this study, the test performed here extended their research to broaden the time points for early-stage sperm storage. Therefore, with the results of their study and the one performed in this study, MHBSS was chosen as the extender for motility, status, viability, and hypoosmotic swelling tests to eliminate any possible influence of poor extender quality on sperm performance.

The viability and hypoosmotic reactivity of sperm were expected to demonstrate similar results, linking viability and membrane integrity. However, viability showed no significant difference between the mutant lines and WT [AB]. HOST showed highly significant changes in reactivity between ALG1 and DPAGT1 mutant lines with WT [AB] indicating the integrity of the spermatozoon’s plasma membrane was not a direct indicator of its viability (Figure 4). In comparison of wild type sperm viability of prior research and the viability measured here, the viability of WT [AB] sperm shown in Figure 4A was much lower than the average range of sperm viability demonstrated by Cattelan et al. (2021). They found a range of viability from approximately 98% to 95% depending on the time between sperm collection. This also demonstrated that the buildup of ROS within the testis was not impactful on wild type sperm even when stored within them for 4 to 12 days (Cattelan et al., 2021). The low percentage of viable cells for mutants may be attributed to an excess of ROS within the testis during storage; although, this cannot account for the low percentage of viable cells for WT [AB] shown here.

Based on HOST results from Cabrita et al. (1999), the WT [AB] would have had approximately 60% to 70% reactive cells based on the viability of 90%. Instead, the reactive cells of WT [AB] fell between 80% to 100%, which further demonstrated the nonlinear relationship of viability and hypoosmotic swelling (Figure 4C). Results of HOST showed highly significant decreases in reactiveness for two of the three mutant lines. The increase of nonreactive, lysed cells was demonstrative of the poor membrane integrity of CDG mutant sperm cells. As mentioned previously, high levels of ROS, such as radical hydroxide, were expected to cause damage to cellular membranes, especially the plasma membrane (Shyr et al., 2025). The decrease in reactivity and the increase in lack of reaction was indicative of this effect. Thus, the results of the HOST suggested that the presence of a CDG likely caused membrane damage, decreasing sperm membrane integrity.

While results met expectations, excluding status and viability, several confounding variables were not apparent in this study. One such variable was the collection of sperm and the inconsistency of productive stripping. Mutant zebrafish were inconsistent in the amount of sperm produced with each stripping procedure. Often, mutants would produce little to no sperm or expressed urine alone. Unlike the mutants, WT [AB] produced similar volumes and densities of sperm with each collection. While the only known difference between the mutant line and WT [AB] zebrafish was the presence of a CDG or CDDG, the inconsistent and irregular collection of sperm from mutants cannot be expressly connected to the mutations alone without further analysis. There was also a small sample size factor for each sperm analysis, excluding the extender comparison test. This was due to the availability of sexually mature and size appropriate males for the mutant lines. Because of lethality for ALG1 and DPAGT1, a quarter of all progenies were always nonviable, and a quarter were always wild type. Half of those surviving were expected to be male, cutting the number of zebrafish produced from each breeding significantly. Then there was the factor of age; all zebrafish used from WT [AB], ALG1, and DPAGT1 lines were approximately 18 months old. NGLY1 zebrafish were approximately 22 months old. These ages were at the upper end of the average zebrafish lifespan. This was done to ensure all fish were around the same age and based on the availability of fish from different age groups. These factors encourage replication or reconstruction of this study and further regard to analysis of influential factors.

Despite the possible influence of confounding variables, results of this study encourage further subcellular research. Since NGLY1 homozygous mutants were not lethal in the zebrafish model, comparison of the heterozygous and homozygous mutant would be a plausible next step in understanding CDDGs. Further insight into the direct driver which affected the decrease in concentration, motility, and hypoosmotic swelling would pilot a better understanding of the mechanisms which negatively impact the cell mutated by enzyme loss of function. This would not only reflect the mechanisms in sperm cells but also somatic cells when addressing the possibility of Sertoli cells influencing sperm quality. Histological and proteomic analyses of sperm cells and Sertoli cells are desirable targets for further research.

## Conclusion

CDGs and CDDGs are congenital disorders arising from cells with genetic mutations which disrupt the processing of glycans. Sperm quality is influenced both by somatic Sertoli cells and developing spermatocytes. A simple collection of sperm cells allows for examination of the disorders’ effect by analyzing parameters of sperm quality. This study finds that CDGs and CDDGs may decrease sperm concentration, motility, and hypoosmotic swelling by possibly inducing cellular stress and damaging the plasma membrane. Further research is necessary to determine the specific driver in decreasing sperm quality and to better understand the mechanisms these disorders cause harm.

